# Targeted sequencing of venom genes from cone snail genomes reveals coupling between dietary breadth and conotoxin diversity

**DOI:** 10.1101/107672

**Authors:** Mark A Phuong, Gusti N Mahardika

**Author notes:** Corresponding author: M. A. Phuong.

## Abstract

Although venomous taxa provide an attractive system to study the genetic basis of adaptation and speciation, the slow pace of toxin gene discovery through traditional laboratory techniques (e.g., cDNA cloning) have limited their utility in the study of ecology and evolution. Here, we applied targeted sequencing techniques to selectively recover venom gene superfamilies and non-toxin loci from the genomes of 32 species of cone snails (family, Conidae), a hyper diverse group of carnivorous marine gastropods that capture their prey using a cocktail of neurotoxic proteins (conotoxins). We were able to successfully recover conotoxin gene superfamilies across all species sequenced in this study with high confidence (> 100X coverage). We found that conotoxin gene superfamilies are composed of 1-6 exons and adjacent noncoding regions are not enriched for simple repetitive elements. Additionally, we provided further evidence for several genetic factors shaping venom composition in cone snails, including positive selection, extensive gene turnover, expression regulation, and potentially, presence-absence variation. Using comparative phylogenetic methods, we found that while diet specificity did not predict patterns of conotoxin gene superfamily size evolution, dietary breadth was positively correlated with total conotoxin gene diversity. These results continue to emphasize the importance of dietary breadth in shaping venom evolution, an underappreciated ecological correlate in venom biology. Finally, the targeted sequencing technique demonstrated here has the potential to radically increase the pace at which venom gene families are sequenced and studied, reshaping our ability to understand the impact of genetic changes on ecologically relevant phenotypes and subsequent diversification.

## Introduction

Understanding the molecular basis for adaptation and speciation is a central goal in evolutionary biology. Past studies have described several genetic characteristics that seem to be associated with rapidly radiating clades or the evolution of novel phenotypes, including evidence for diversifying selection, gene gains and losses, and accelerated rates of sequence evolution (Henrissat et al. 2012; Guillén et al. 2014; Brawand et al. 2014; Cornetti et al. 2015; Malmstrøm et al. 2016; Pease et al. 2016). Although large-scale comparative genomic studies have vastly increased our knowledge of the genetic changes associated with diversification, the link between genotype and ecologically relevant phenotypes frequently remains unclear. Often, the functional consequences of genetic patterns such as an excess of gene duplicates or regions under positive selection are unknown (Brawand et al. 2014; Cornetti et al. 2015; Pease et al. 2016), limiting our ability to understand how genetic changes shape the evolutionary trajectory of species.

Animal venoms provide an excellent opportunity to study the interplay between genetics and adaptation because of the relatively simple relationship between genotype, phenotype, and ecology. Venoms have evolved multiple times throughout the tree of life (e.g., spiders, snakes, and snails) and play a direct role in prey capture and survival (Barlow et al. 2009; Casewell et al. 2013). Venoms are composed of mixtures of toxic proteins and peptides that are usually encoded directly by a handful of known gene families (Kordis & Gubensek 2000; Fry et al. 2009; Casewell et al. 2013). Exceptionally high estimated rates of gene duplication and diversifying selection across these venom genes families are thought to contribute to the evolution of novel proteins and thus changes in venom composition (Duda & Palumbi 1999; Gibbs & Rossiter 2008; Chang & Duda 2012), allowing venomous taxa to specialize and adapt onto different prey species (Kohn 1959a; Daltry et al. 1996; Li et al. 2005; Barlow et al. 2009; Chang & Duda 2016; Phuong et al. 2016). Therefore, the study of venomous taxa can facilitate understanding of the genetic contributions to ecologically relevant traits and subsequent diversification.

A fundamental challenge associated with the study of venom evolution is the inability to rapidly obtain sequences from venomous multi-gene families. Traditionally, venom genes were sequenced through cDNA cloning techniques, which can be labor intensive and time-consuming (Gibbs & Rossiter 2008; Chang et al. 2015). Although transcriptome sequencing has greatly increased the pace of venom gene discovery (e.g., Casewell et al. 2009; Phuong et al. 2016), transcriptome sequencing still requires fresh RNA extracts from venom organs, which may be difficult to obtain for rare and/or dangerous species. Targeted sequencing approaches have vastly improved the capacity to obtain thousands of markers across populations and species for ecological and evolutionary studies (Faircloth et al. 2012; Bi et al. 2012). Until now, these approaches have not been applied to selectively sequence venomous genomic regions. This may be in part, due to the extraordinary levels of sequence divergence exhibited by venom loci (Gibbs & Rossiter 2008; Chang & Duda 2012), potentially rendering probes designed from a single sequence from one gene family unable to recover any other sequences in the same family (Fig. 1). However, past studies have shown that noncoding regions adjacent to hypervariable mature toxin exons are conserved between species (Nakashima et al. 1993, 1995; Gibbs & Rossiter 2008; Wu et al. 2013), suggesting that these conserved regions can be used for probe design to potentially recover all venom genes across clades of venomous taxa.

**Figure 1.**
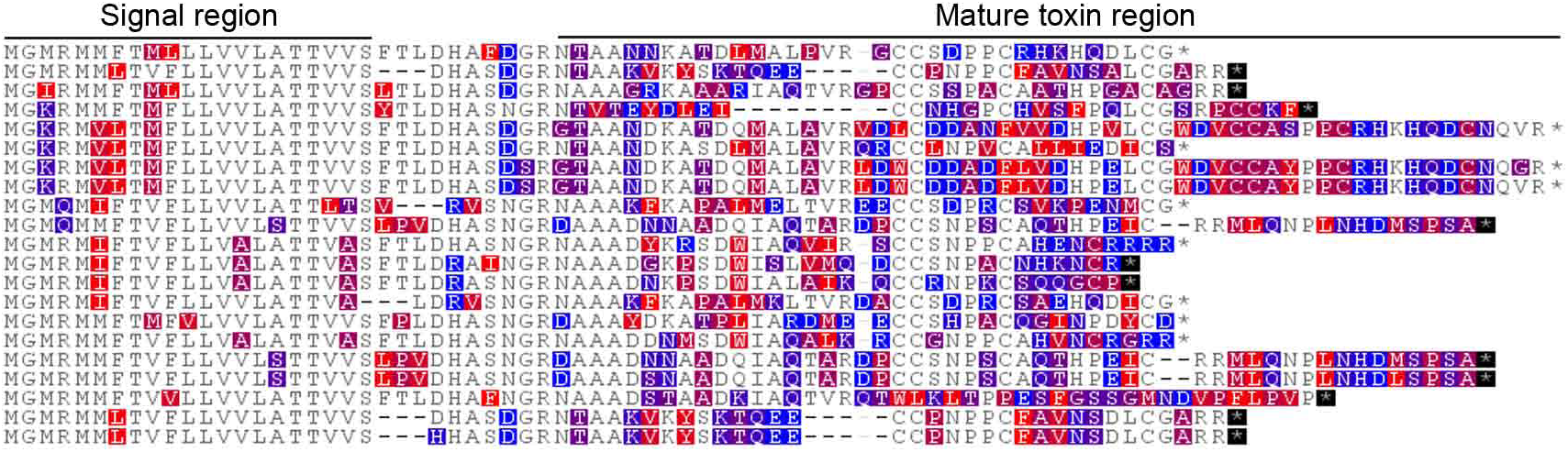
A superfamily conotoxins from *Conus lividus* described in the transcriptome from Phuong et al. 2016.

Here, we used a targeted sequencing approach to recover venom genes and study the evolution of venom gene families across 32 species of cone snails from the family, Conidae. Cone snails are a hyper diverse group of carnivorous marine gastropods (> 700 spp.) that capture their prey using a cocktail of venomous neurotoxins (Puillandre et al. 2014). Cone snail venom precursor peptides (conotoxins) are typically composed of three regions: the signal region that directs the protein into the secretory pathway, the prepro region that is cleaved during protein maturation, and the mature region that ultimately becomes the mature peptide (Robinson & Norton 2014). In some instances, there exists a ‘post’ region of the peptide following the mature region that is also cleaved during protein processing (Robinson & Norton 2014). Conotoxins are classified into > 40 gene superfamilies (e.g., A superfamily, O1 superfamily, etc.) based on signal sequence identity, though some gene superfamilies were identified based on domain similarities to proteins from other venomous taxa (Robinson & Norton 2014). To examine the evolution of conotoxin gene superfamilies from genomic DNA, we designed probes targeting over 800 non-conotoxin genes for phylogenetic analyses and conotoxins from 12 previously sequenced Conidae transcriptomes (Phuong et al. 2016). With the recovered conotoxin loci, we describe several features of conotoxin genes, including its genetic architecture, molecular evolution, expression patterns, and changes in gene superfamily size. Finally, we use comparative phylogenetic methods to test whether diet specificity or dietary breadth can explain patterns of gene superfamily size evolution.

## Results

### Exon capture results

We used custom designed 120bp baits (MYbaits; Mycroarray, Ann Arbor, Michigan, USA) to selectively target phylogenetic markers and conotoxin genes from 32 Conidae species (Table S1). We sequenced all samples on a single Illumina HiSeq2000 lane, producing an average of 12.8 million reads per sample (Table S1). After redefining exon boundaries for the phylogenetic markers, we generated a reference that consisted of 5883 loci. We recovered an average of 5335 loci (90.7%) across all samples representing ~0.66 Mb (Megabases) on average (Table S1). For the conotoxin genes, the number of gene models we assembled containing a conotoxin exon ranged from 281 fragments in *Conus papilliferus* to 2278 fragments in *C. atristophanes* (Table S1). Approximately 48.8% of the reads mapped to both the phylogenetic markers and venom genes with 52.3% of these reads being marked as duplicates (Table S1). Average coverage across the phylogenetic markers was 95.9X, while the average coverage for the conotoxin exons was 149.6X (Table S1).

We recovered representative exons from all 49 conotoxin gene superfamilies targeted, plus exons from the Q gene superfamily which we did not explicitly target (Fig. S1). Of the 49 targeted gene superfamilies, ‘capture success’ (defined in Materials and Methods) was 80% or above for 34 gene superfamilies, even though we did not explicitly target every single transcript (Table S2). For example, we only targeted 1 sequence of the A gene superfamily from *C. varius*, but we recovered sequences that showed high identity to every single transcript from the A gene superfamily discovered in the *C. varius* transcriptome (Table S2). We assessed the ability of targeted sequencing to recover conotoxins from species that were not explicitly targeted in the bait sequences by calculating the number of previously sequenced conotoxins (obtained via Genbank and Conoserver (Kaas et al. 2010)) recovered in our dataset. The number of previously sequenced conotoxins from online databases recovered in this study did not differ depending on whether or not a species was explicitly targeted by our bait design (t-test, t = -0.8, df = 20.1, p-value > 0.05, Table S3).

### Conotoxin genetic architecture

Through analyses of conotoxin genetic structure across species, we found that the number of exons that comprise a conotoxin transcript ranged from 1 to 6 exons and exon size ranged from 5bp to 444bp (Table S4). Whether or not untranslated regions (UTRs) were adjacent to terminal exons was dependent on the gene superfamily, with some gene superfamilies always having both 5’ and 3’ UTRs adjacent to terminal exons and some where the 5’ or 3’ UTRs cannot be found directly adjacent to the terminal exons (Table S4). Regions in conotoxin transcripts identified as the signal region, the mature region, or the post region were most often confined to a single exon (Fig. 2). In contrast, the prepro region was more frequently distributed across more than one exon (Fig. 2).

**Figure 2.**
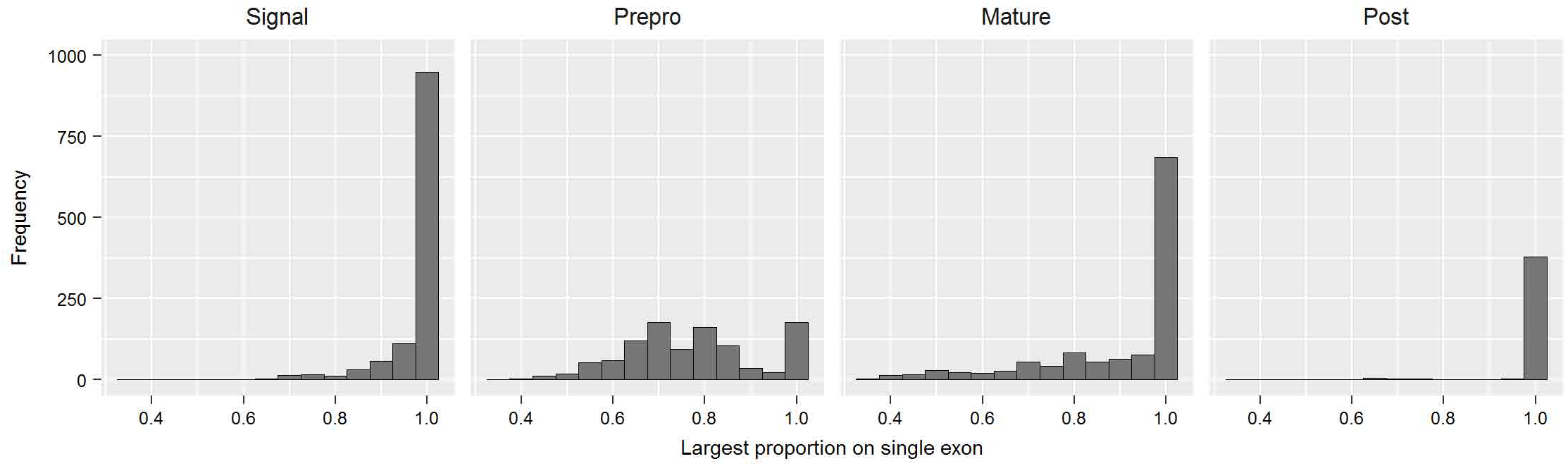
Histograms showing the frequency of the largest proportion of conotoxin precursor peptide regions found on a single exon in Conidae genomes.

Because preliminary studies on conotoxin genetic structure suggested that conotoxin introns are enriched for simple repeats (Wu et al. 2013; Barghi et al. 2015a), we assessed simple repeat content in noncoding regions from both conotoxin and non-conotoxin loci. The percentage of contigs containing a simple repeat in noncoding regions adjacent to conotoxin exons (avg = 52.8%) was lower compared to non-conotoxin loci (avg = 55.6%, Table S5). The percentage length of simple repeats in noncoding regions near conotoxins (avg = 2.9%) was higher than in non-conotoxin loci (avg = 2.6%, Table S5). While paired t-tests found significant differences in the means in both measures of simple repeat abundance, the differences of the means were quite small in both the percentage of contigs containing simple repeats in noncoding regions (diff = 2.8%, t = -3.05, df = 31, p-value = 0.005) and the percentage length of simple repeats in these regions (diff = 0.3%, t = 4.53, df = 31, p-value < 0.0001).

### Conotoxin molecular evolution

To determine if there are differences in divergence depending on what conotoxin precursor peptide region each exon contains, we quantified the level of sequence divergence between exons and immediately adjacent noncoding regions. Exons containing the signal region were more conserved than their adjacent regions (average relative ratio < 1, Table S6, Fig. 3, S2). In contrast, all other exon classifications generally showed the opposite pattern, where the exons were typically more divergent relative to their adjacent noncoding regions (average relative ratio > 1, Table S6, Fig. 3, S2). The largest contrast in divergence between exons and adjacent noncoding regions came from exons containing the mature region, where the coding region was on average 2.9 times more divergent than regions surrounding the exon (Table S6, Fig. 3, S2). For comparison, exons from non-conotoxin genes were more conserved than their adjacent regions (relative exon to adjacent region divergence ratio < 1, Fig. 3).

**Figure 3.**
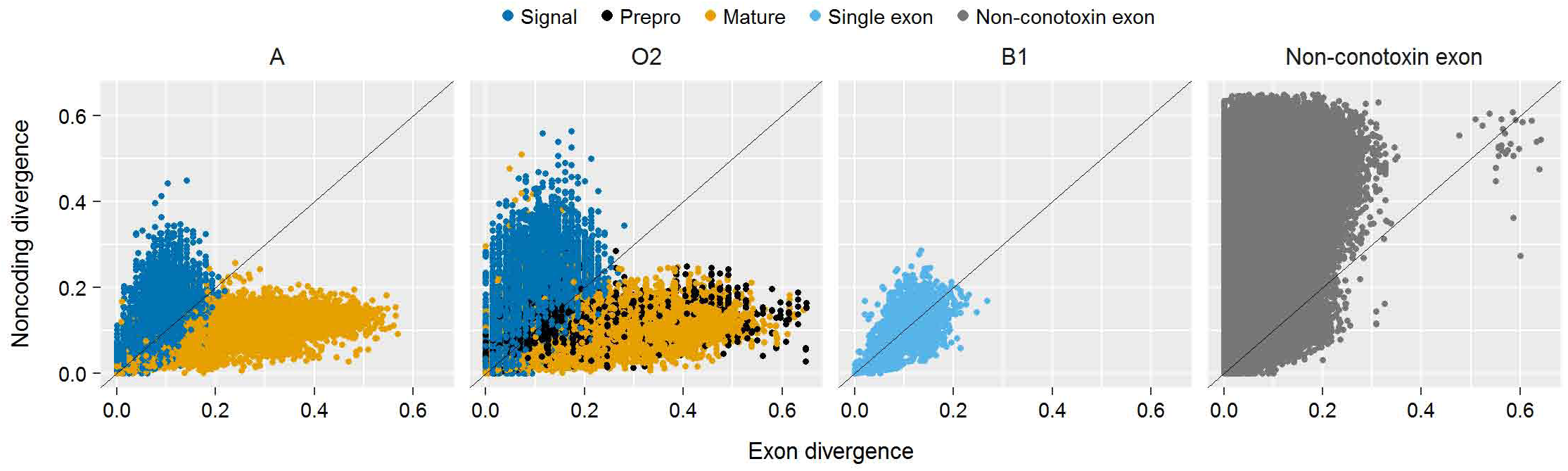
Scatterplot of uncorrected pairwise distances for select gene superfamilies and non-conotoxin loci between exons and adjacent noncoding regions.

### Conotoxin expression

The proportion of conotoxin genes expressed per gene superfamily was highly variable (Table S7) and the exact proportion depended on the gene superfamily and the species. In several cases, all gene copies of a gene superfamily were not expressed in the transcriptome (e.g., *Conus ebraeus,* A gene superfamily, 0/9 copies expressed, Table S7), and in other cases, all copies were expressed in the transcriptome (e.g., *C. californicus*, O3 gene superfamily, 3/3 copies expressed, Table S7). The average proportion of gene copies expressed per gene superfamily per species was 45% (range: 24% – 63%, Table S7).

### Conotoxin gene superfamily size evolution

With a concatenated alignment of 4441 exons representing 573854bp, we recovered a highly supported phylogeny with all but 4 nodes having ≤ 95% bootstrap support (Fig. 4). Total conotoxin gene diversity ranged from as low as 120 in *C. papilliferus* to as high as 859 in *C. coronatus* (Fig. 4). 25 gene superfamilies showed evidence of phylogenetic signal in gene superfamily size, such that closely related species tended to have similar gene superfamily sizes (Table S8). For example, nearly all of the gene copies from the A superfamily in a clade consisting of *C. mustelinus, C. capitaneus, C. miles, C. vexillum*, and *C. rattus* seemed to have been lost from the genome, while a clade composed of *C. lischkeanus, C. muriculatus,* and *C. lividus* seemed to have expanded in membership size (Fig. 4). CAFE (Han et al. 2013) estimates of net gene gains and losses showed that species-specific net conotoxin expansions and contractions are scattered throughout the phylogeny (Fig. 4, Fig. S3).

**Figure 4.**
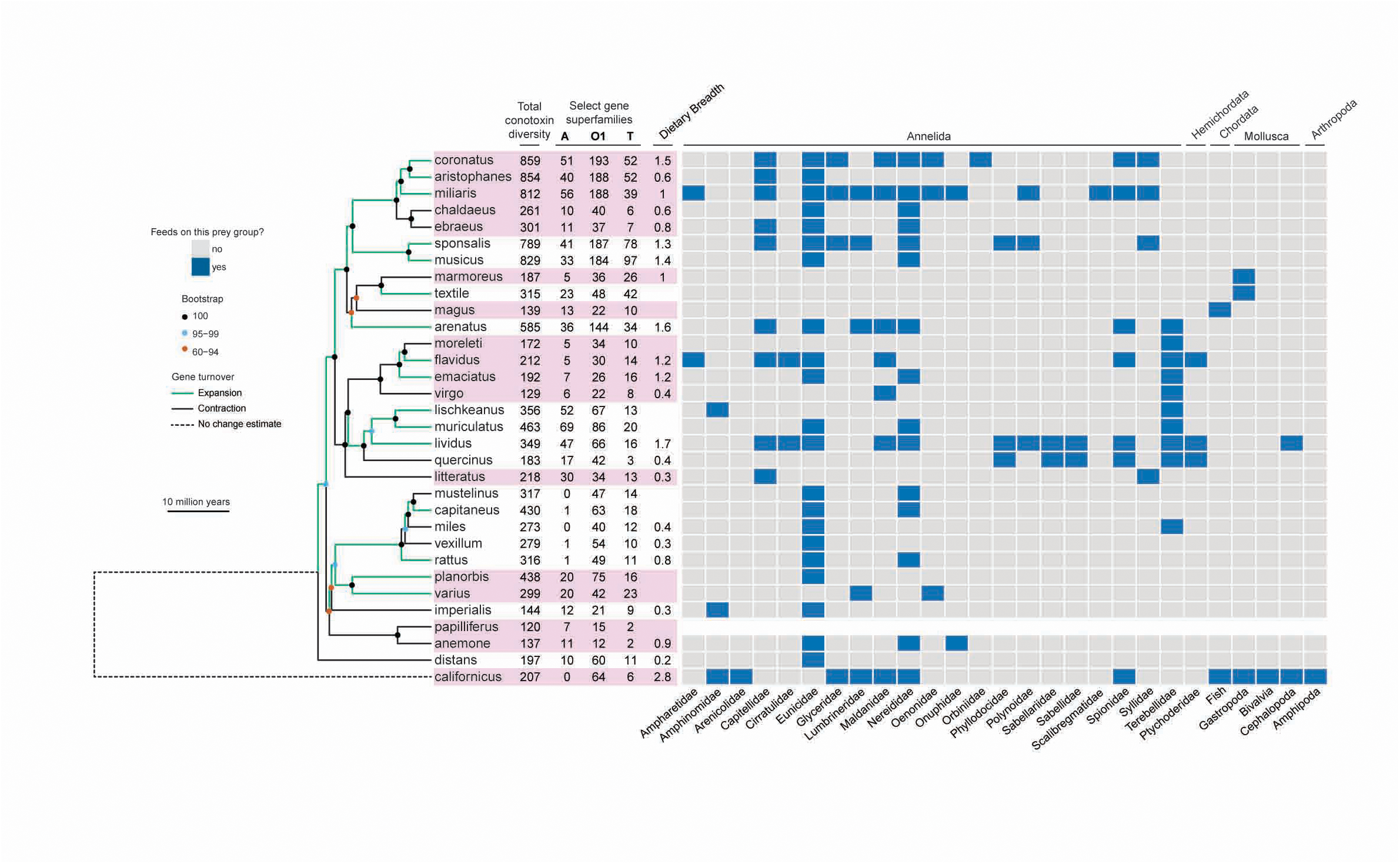
Diet and conotoxin evolution in a phylogenetic context. Time-calibrated maximumlikelihood phylogeny of 32 Conidae species generated from concatenated alignment of 4441 exons. Total conotoxin diversity and size estimates for commonly studied gene superfamilies displayed next to tip names. Branches are colored base on net gains or losses in total conotoxin diversity based on CAFE analyses. Recognized subgenera are alternately colored pink.

### Diet and conotoxin gene superfamily evolution

We used comparative phylogenetic methods and extensive prey information from the literature to examine the impact of diet specificity (i.e. what prey a cone snail feeds upon) and dietary breadth (i.e., how many prey species a cone snail feeds upon) on the evolution of conotoxin gene diversity. Neither diet specificity nor dietary breadth was correlated in changes with gene superfamily size (D-PGLS [distance-based phylogenetic generalized least squares], p > 0.05). While diet specificity did not predict changes in total conotoxin diversity (PGLS, p > 0.05), we found a significant positive relationship between dietary breadth and total conotoxin diversity in both the full conotoxin dataset (PGLS, p < 0.05, Fig. 5) and the conotoxin dataset containing gene superfamilies that had > 80% capture success (PGLS, p < 0.001).

**Figure 5.**
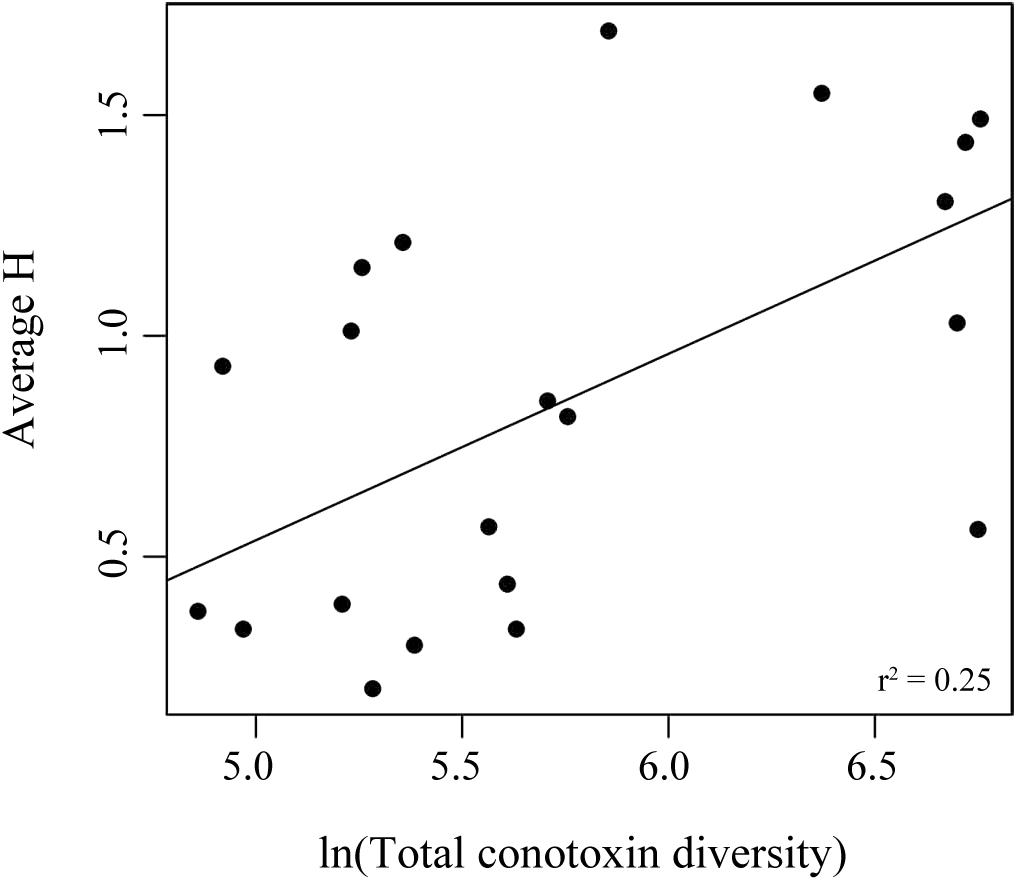
Scatterplot of total conotoxin gene diversity and dietary breadth. Graph labeled with correlation coefficient.

## Discussion

### Targeted sequencing and conotoxin discovery

Through targeted sequencing of conotoxins in cone snails, we demonstrate the potential to rapidly obtain venom sequences at high coverage (> 100X, Table S1) from species for which no venom information is available and without the need of RNA from the venom duct. This is remarkable, given that alignments in amino acid sequences between mature regions of a single gene superfamily within a single individual can be incomprehensible (Fig. 1) due to the high evolutionary rates of the mature region (Duda & Palumbi 1999). Effective capture of conotoxin gene superfamilies was possible in part because conotoxin exons were often directly adjacent to conserved UTRs, which were targeted in the design (Table S4). For several gene superfamilies, exons containing the prepro region or the mature region were not immediately adjacent to the UTRs – thus, these gene superfamilies were not as effectively recovered from species not explicitly targeted by the baits (e.g., prepro region, gene superfamily O3, Fig. S1). However, we recovered intron sequences in this study that can be used in future bait designs to anchor probes to effectively recover the entire coding region of a conotoxin transcript because adjacent noncoding regions are often evolving at a much slower rate than the coding region (Table S6, Fig. 3, S2). While it has been recognized for decades that cone snails collectively harbor tens of thousands of biologically relevant proteins for fields such as molecular biology and pharmacology in their venoms (Olivera & Teichert 2007; Lewis 2009), traditional techniques for conotoxin sequencing (e.g., cDNA cloning) have barely begun to uncover and characterize the full breadth of conotoxin diversity. The targeted sequencing technique presented in this study has the potential to fundamentally reshape the speed at which conotoxins are discovered, potentially having significant implications for the study of conotoxins in molecular biology, pharmacology, ecology, and evolution.

When compared to conotoxin sequences available on Genbank and ConoServer, we found that we were able to recover approximately the same proportion of previously sequenced conotoxins across species regardless of whether the species was explicitly targeted with the baits (Table S3). We performed a coarse investigation of database conotoxins and determined potential reasons for why we were not able to recover certain previously sequenced conotoxins. These reasons include: (a) the species in the database was misidentified, (b) the database conotoxin had no reliable reference in the literature, (c) the conotoxin was present in our species, but we could not recover it with the current bait design, and (d) the conotoxin was recoverable (i.e., high sequence similarity to bait sequences designed in this study), but the gene was not present in the genome of the individual used in this study, indicative of presence-absence variation shaping the venom repertoire of cone snail species. Future studies studying the genomic content of conotoxins from the genome will enable understanding of the contributions of presence-absence variation to the evolution of cone snail venom composition.

### Genetic characteristics of conotoxins

Previous studies noted the enrichment of simple repetitive elements (e.g., ATATATAT) in noncoding regions adjacent to conotoxin loci, suggesting that these repetitive regions may be important for conotoxin gene splicing and the regulation of gene expression (Wu et al. 2013; Barghi et al. 2015a). Although our comparisons of simple repeat content between conotoxin and non-conotoxin exons showed that a higher proportion of contigs contained simple repeats near non-conotoxin exons and simple repeats constituted a greater proportion of noncoding length near conotoxin exons, these differences were quite small (Table S5), suggesting that the presence of simple repeats are not unique to conotoxins. This discrepancy occurred between this study and past work because previous studies did not compare repeat content to non-conotoxin loci (Wu et al. 2013; Barghi et al. 2015a), underscoring the importance of making appropriate comparisons to identify features that may be exclusive to conotoxin evolution.

The ratio of exon to noncoding divergence depended on what conotoxin region was encoded by the exon, aligning with previous work characterizing rate variation in snake venom proteins (Nakashima et al. 1993, 1995; Gibbs & Rossiter 2008). Specifically, the exon containing the signal region was conserved and evolved much more slowly than adjacent noncoding regions (Table S6, Fig. 3, S2). This is similar to the pattern found in non-conotoxin exons (Fig. 3), indicative of purifying selection removing deleterious mutations from coding regions of critically important proteins (Hughes & Yeager 1997). In contrast, the exon diverges faster than the noncoding regions in all other exons, with the clearest difference between exon and noncoding region divergence seen in the exon containing most or all of the mature toxin region. This same pattern is also seen in other genes under positive selection, such as PLA2 genes in snakes (Nakashima et al. 1993, 1995; Gibbs & Rossiter 2008) and fertilization genes in abalone (Metz et al. 1998). Although we did not use traditional methods to test for positive selection (e.g., MK tests, etc.), positive selection is well documented in cone snails (Duda & Remigio 2008; Duda 2008; Puillandre et al. 2010) and is therefore inferred to shape patterns of increased divergence in coding regions relative to noncoding regions.

We found that on average, cone snails only express a fraction of the conotoxin genes available in their genomes, concurring with similar reports from smaller sets of gene superfamilies (Chang & Duda 2012, 2014; Barghi et al. 2015a). Several reasons could lead to this pattern. First, it is known that expression changes throughout an individual’s lifetime in cone snails (Barghi et al. 2015b; Chang & Duda 2016), suggesting that the complement of genes expressed in the transcriptomes from Phuong et al. 2016 represent the adult conotoxins, and genes not discovered in the transcriptome but recovered from the genome are genes that are expressed in other life stages. Second, prey taxa available to cone snail species change with geography and so do the conspecifics it must compete against (Kohn 1959a, 1978; Kohn & Nybakken 1975; Duda & Lee 2009; Chang et al. 2015); therefore, different genes may be turned on or off in different geographic localities depending on the prey resources available and the composition of competitors in an individual’s environment. Finally, some of the conotoxin genes in the genome may not be expressed because they are no longer functional and have become psuedogenized. Future work comparing patterns of expression relative to genomic availability will be able to disentangle the impact of conotoxin expression on changes to the venom phenotype.

We detected evidence for phylogenetic signal in the membership size of 25 gene superfamilies (Table S8, Fig. 4), suggesting that history plays a role in shaping gene gains and losses in cone snails. We note that uncovering evidence for phylogenetic signal in gene superfamily size does not imply that natural selection has not played a role in the evolution of venom as implied in (Gibbs et al. 2013). As described in Revell et al. (2008), evolutionary processes should not be inferred from patterns of phylogenetic signal because several contrasting models of trait evolution can lead to similar amounts of phylogenetic signal. Through CAFE analyses, we also showed that venom composition is shaped by both net gains and losses in the entire genomic content of conotoxins (Fig. 4, Fig. S3). This result is in line with past studies showing that gene turnover is a fundamental characteristic shaping species’ genic venom content (Duda & Palumbi 1999; Chang & Duda 2012; Dowell et al. 2016).

### Diet and venom evolution

Why do cone snails vary in conotoxin gene superfamily size? Contrary to the popular assumption that particular gene superfamilies are associated with certain prey items (e.g, Kaas et al. 2010; Jin et al. 2013), diet categories did not predict changes in gene superfamily size or total conotoxin diversity. This result aligns with a growing body of literature suggesting that the specific prey a species feeds upon may not accurately predict patterns of conotoxin evolution (Puillandre et al. 2012; Chang et al. 2015; Phuong et al. 2016). While dietary breadth also did not predict changes in gene superfamily size, we found a significant positive relationship with total conotoxin diversity (Fig. 5), aligning with several studies showing a tight coupling between dietary breadth and venom gene diversity in cone snails at nearly all biological scales of organization (Duda & Lee 2009; Chang et al. 2015; Chang & Duda 2016; Phuong et al. 2016). The importance of dietary breadth shaping venom evolution remains underappreciated and untested in other venomous systems despite strong signals across several studies in cone snails. Future work examining the role of dietary breadth in shaping the evolution of venom in other venomous taxa will greatly advance our understanding between the interplay between diet and venom. The lack of a relationship between dietary breadth and changes in conotoxin gene superfamily size suggests that venom should be characterized as an aggregate trait rather than decomposed into individual parts to fully assess the impact of dietary breadth on conotoxin evolution. Indeed, conotoxin gene superfamilies provide little information on protein function and it is known that proteins with similar functions can evolve convergently in different gene superfamilies (Kaas et al. 2010; Puillandre et al. 2012; Robinson & Norton 2014). Further, studies have documented synergistic and complementary effects of conotoxins on prey species, suggesting that selection may act on the entire cocktail rather than individual components (Olivera 1997).

## Conclusions

Through targeted sequencing of conotoxin genes, we provide additional evidence for several genetic characteristics that shape venom composition in cone snails, including presence-absence variation, positive selection, regulation of gene expression, and gene turnover. In addition, we find that variation in conotoxin diversity tracks changes in dietary breadth, suggesting that species with more generalist diets contain a greater number of conotoxin genes in their genome. Given that increased gene diversity is thought to confer an increased capacity for evolutionary change and species diversification (Kirschner & Gerhart 1998; Yang 2001; Malmstrøm et al. 2016), generalist species may speciate at faster rates than species with specialist diets. The targeted sequencing technique presented in this paper provides the necessary methodological advancement to rapidly sequence toxin genes across diverse clades of species, allowing tests of the relationship between ecology, toxin gene diversity, and higher order biodiversity patterns to be realized in future work.

## Materials and Methods

### Bait design and data collection

To recover markers for phylogenetic analyses, we targeted 886 protein coding genes representing 728,860bp. 482 of these genes were identified to be orthologous in Pulmonate gastropods (Teasdale et al. 2016) and we identified the remaining 404 genes using a reciprocal blast approach with 12 Conidae transcriptomes from (Phuong et al. 2016). For each gene, we chose the longest sequence from one of the 12 Conidae transcriptomes as the target sequence. For 421 of these genes, we used the entire length of the sequence as the target sequence, while for the remaining genes, we sliced the target sequences into smaller components based on exon/intron boundaries inferred with EXONERATE (Slater & Birney 2005) using the *Lottia gigantea* genome as our reference. If exons were below 120bp (i.e., our desired bait length) in length, but longer than 50bp, we generated chimeric target sequences by fusing immediately adjacent exons. We tiled bait sequences every 60bp across each target sequence. For the conotoxin genes, we targeted 1147 conotoxins discovered from an early analysis of the 12 transcriptomes described in (Phuong et al. 2016). These sequences represent regions targeting 49 gene superfamilies and we included the full protein coding region plus 100bp of the 5’ and 3’ untranslated regions in our bait design when possible (Table S2). We tiled bait sequences every 40 bp across each conotoxin target sequence.

We obtained tissue samples for 32 Conidae species through field collections in Australia and Indonesia and from the collections at the Australian Museum in Sydney, Australia (Table S1). We extracted genomic DNA using an EZNA Mollusc DNA kit (Omega Bio-Tek, Doraville, GA, USA) and prepared index-specific libraries following the Meyer and Kircher (2010) protocol. We pooled 8 samples at a total concentration of 1.6μg DNA per capture reaction and allowed the baits to hybridize with the DNA for ~24 hours. We substituted the universal blockers provided with the MYbaits kit with xGEN blockers (Integrated DNA Technologies). After hybridization, we sequenced all 32 samples on a single lane HiSeq2000 lane with 100bp paired-end reads.

### Data assembly, processing, and filtration

We trimmed reads for quality and adapter contamination using Trimmomatic (Bolger et al. 2014), merged reads using FLASH (Magoč & Salzberg 2011), generated assemblies for each sample using SPAdes (Bankevich et al. 2012) and reduced redundancy in the assemblies with cap3 (Huang & Madan 1999) and cd-hit (Li & Godzik 2006).

For the phylogenetic markers, we used BLAST+ (Altschup et al. 1990) to associate assembled contigs with the target sequences and used EXONERATE to redefine exon/intron boundaries because either (a) exon/intron boundaries were never denoted or (b) previously defined exons were composed of smaller fragments. For each sample, we used bowtie2 (Langmead & Salzberg 2012) to map reads to a reference containing only the contigs associated with the original target sequences and marked duplicates using picard-tools (http://broadinstitute.github.io/picard). We masked all positions that were below 5X coverage and removed the entire sequence if > 30% of the sequence was masked. To filter potential paralogous sequences in each species, we calculated heterozygosity (number of heterozygous sites/total number of sites) for each locus by identifying heterozygous positions using samtools and bcftools (Li et al. 2009) and removed loci that were at least two standard deviations away from the mean heterozygosity.

For the conotoxin sequences, it is known that traditional assemblers perform poorly in reconstructing all potential conotoxin gene copies (Lavergne et al. 2015; Phuong et al. 2016). To ameliorate this issue, we reassembled conotoxin genes using the assembler PRICE (Ruby et al. 2013), which employs iterative mapping and extension using paired read information to build out contigs from initial seed sequences. To identify potential seed sequences for contig extension, we first mapped reads to the entire assembly outputted by SPAdes using bowtie2. Then, we identified all sequence regions that locally aligned to any part of the original conotoxin target sequences via blastn; these regions represented our preliminary seed sequences. We kept all preliminary seed sequences that were at least 100 bp (read length of samples in this study) and extended these seeds to 100bp if the alignable region was below that threshold. When extending these initial regions, we used Tandem Repeats Finder (Benson 1999) to identify simple repeats and minimize the presence of these genomic elements in the preliminary seed sequences. Often, only a subset of conotoxins are fully assembled with traditional assemblers (Phuong et al. 2016). However, when reads are mapped to these assemblies, unique conotoxin loci are similar enough to each other that relaxed mapping parameters will allow multiple copies to map to the contigs that were assembled. Therefore, multiple conotoxin copies will often map to each preliminary seed sequence. To generate seed sequences for all unique conotoxin loci, we used the python module pysam (https://github.com/pysam-developers/pysam) to pull all reads that mapped to regions of contigs representing the preliminary seed sequence and we reconstructed contigs from these reads using cd-hit and cap3. From these reconstructed contigs, we used blastn to identify >100bp regions that matched the original preliminary seed and used these hits as our final seeds. We merged all final seeds that were 100% identical using cd-hit, mapped reads to these seeds using bowtie2, and used PRICE to re-assemble and extend each seed sequence under 5 MPI values (90%, 92%, 94%, 96%, 98%) with only the set of reads that mapped to that initial seed. A sequence was successfully reassembled if it shared ≥ 90% identity to the original seed sequence and if the final sequence was longer than the initial seed. For each seed sequence, we only retained the longest sequence out of the 5 MPI iterations for downstream filtering.

In order to generate a conotoxin reference database containing sequences that included both exons and adjacent noncoding regions, we used blastn and exonerate on species that were used in the bait design to (a) perform species-specific searches between our reassembled contigs and a reference containing all conotoxins in (Phuong et al. 2016) and (b) define exon/intron boundaries on our reassembled contigs. In cases where a predicted terminal exon (i.e., the first or last exon of a conotoxin) was short (< 40 bp) and did not blast to any reassembled contig in our exon capture dataset, we replaced the reference conotoxin from Phuong et al 2016 with a conotoxin containing the adjacent UTR regions to aid in the sequence searches. We generated conotoxins with UTR regions using the PRICE algorithm as described above because the reference conotoxins from the final dataset in Phuong et al. 2016 did not include the UTR regions. With this final conotoxin reference containing sequences with exons and introns pre-defined, we used blastn to associate contigs with this reference in every species and used exonerate, blastx, and tblastn to define exon/intron boundaries. When exon/intron boundaries could not be defined through these methods, we guessed the boundaries by aligning the assembled contig to the reference sequence using MAFFT and denoted the boundaries across the region of overlap with the exon in the reference sequence. For each sample, we mapped reads using bowtie2, accounted for duplicates using picard-tools, and retained sequences that had at least 10x coverage across the exons defined within each contig. We masked regions below 10x coverage and used cd-hit to merge contigs that were ≥ 98% similar, generating our final conotoxin gene models. Finally, we used HapCUT (Bansal & Bafna 2008) to generate all unique haplotypes across coding regions.

To assess the overall effectiveness of our targeted sequencing experiment, we calculated (a) percent of reads aligned to intended targets, (b) percent duplicates, and (c) average coverage across targeted regions. To assess capture success of conotoxins, we divided the number of conotoxin transcripts successfully recovered in the exon capture dataset by the number of conotoxin transcripts discovered in Phuong et al. 2016 for each gene superfamily. We defined a conotoxin transcript to be successfully sequenced if > 80% of the transcript was recovered in the exon capture experiment with > 90% identity. To assess the ability of targeted sequencing to recover gene superfamily sequences from species that were not explicitly targeted in the bait sequences, we calculated the number of previously sequenced conotoxins that match contigs recovered in our dataset. We gathered conotoxin sequences from Genbank and ConoServer (Kaas et al. 2010) with species names that correspond to species in this study, merged sequences with 98% identity using cdhit, and used blastn to perform species-specific searches. We defined a conotoxin as successfully sequenced if the hypervariable mature toxin coding region aligned with ≥ 95% identity to a sequence in our dataset. We used an unpaired t-test to determine whether the probability of recovering a previously sequenced conotoxin differed depending on whether or not that species’ conotoxin repertoire was incorporated in the bait sequences.

### Conotoxin genetic architecture

To characterize conotoxin genetic architecture, we quantified the following values: (a) the number of exons comprising a conotoxin transcript, (b) average length of each exon, and (c) the size range of exon length. We also determined the proportion of terminal exons adjacent to UTRs by conducting sequence searches (via blastn) between contigs containing terminal exons against a database of conotoxins from Phuong et al. 2016 that were reassembled to contain the UTRs. To determine how traditional conotoxin precursor peptide regions are distributed among exons, we calculated the average proportion of each conotoxin region found on each exon in every gene superfamily. We defined regions of the Phuong et al. 2016 transcripts using ConoPrec (Kaas et al. 2012). We restricted these conotoxin genetic structure analyses to transcripts from Phuong et al. 2016 that were successfully recovered in the exon capture dataset and that were retained after clustering with cd-hit (similarity threshold = 98%). We performed clustering to avoid over-inflating estimates because unique transcripts from Phuong et al. (2016) may have originated from the same gene.

To determine whether conotoxin loci are enriched for simple repetitive elements, we quantified the amount of simple repeats in noncoding regions adjacent to conotoxin exons vs. noncoding regions adjacent to non-conotoxin exons. If multiple exons were found on the same contig, we split the sequences at the midpoint in the intron region to standardize the data because most contigs contain only one exon. We used Tandem Repeat Finder to identify repeats and retained results when (a) the repeat pattern size was between 1 to 6 (i.e., mono-nucleotide repeats to hexa-nucleotide repeats), (b) the pattern was repeated at least 5 times, and (c) patterns within a repeat shared 90% identity with each other. We calculated (a) the proportion of contigs containing simple repeats and (b) the proportion of bases that comprised of simple repeats and tested the differences between conotoxins and non-conotoxin loci using paired t-tests.

### Conotoxin molecular evolution

We first classified all exons into conotoxin precursor peptide regions. For species with transcriptome data, we first labeled exons as either the signal region or the mature region by identifying the exons containing the largest proportion of these separate regions. Then, exons between the signal and mature exon were labeled as the prepro exon(s) and exons after the mature region were labeled as the post exons. Gene superfamilies containing only a single exon were denoted as such. We then used blastn to classify sequences without transcriptome data into these conotoxin precursor peptide regions. For each functional category within each gene superfamily, we calculated uncorrected pairwise distances between all possible pairwise comparisons. To avoid spurious alignments, we only considered comparisons within clusters that clustered with cd-hit at an 80% threshold and we excluded comparisons if (a) the alignment length of the two exons was 20% greater than the longer exon, (b) the align-able nocoding region was below 50bp, or (c) the shorter exon’s length was less than 70% of the length of the longer exon. We calculated separate pairwise distance estimates for regions of the alignment that contained the exon and regions of the alignment that contained the noncoding DNA. We excluded region-labeled exons within superfamilies from this analysis that had less than 50 possible comparisons. For comparison, we also calculated pairwise distances between exons and noncoding regions across our phylogenetic markers which represent non-conotoxin exons, filtered with similar criteria described above.

### Conotoxin expression

To characterize variation in expression patterns among species per gene superfamily, we calculated the number of conotoxin genes expressed in species with transcriptome data divided by the number of genes available in the genome. We restricted these analyses to instances where 90% of the unique mature toxins were recovered for a gene superfamily within a species. To estimate gene superfamily size, we used the exon labeled as containing most or all of the mature region. We defined a conotoxin gene as expressed if we retained a blast hit with 95% identity to a unique mature toxin sequence found in the transcriptome.

### Gene superfamily size change estimation

To compare and contrast gene superfamily size changes between species, we used the total number of exons containing most or all of the signal region as our estimate of gene superfamily size because exons containing the signal regions are relatively conserved across species (Robinson & Norton 2014) and thus have the highest confidence of being recovered through exon capture techniques. To quantify and test the amount of phylogenetic signal in conotoxin gene diversity, we estimated Pagel’s lambda (Pagel 1997) in the R package phytools (Revell 2012). Lambda values range from 0 (phylogenetic independence) to 1 (phylogenetic signal) and p-values < 0.05 represent significant departure from a model of random trait distribution across species with respect to phylogeny. To estimate conotoxin gene superfamily gains and losses along every branch, we used the program CAFEv3.1 (Han et al. 2013), which uses a stochastic gene birth-death process to model the evolution of gene family size. As input, we used a time-calibrated phylogeny and estimates of gene superfamily size for 37 superfamilies that were present in at least 2 taxa. To estimate a time-calibrated phylogeny, we aligned loci that had at least 26 species using MAFFT (Katoh et al. 2005) and used a concatenated alignment to build a phylogeny in RAxML under a GTRGAMMA model of sequence evolution (Stamatakis 2006). We time-calibrated the phylogeny with the program r8s (Sanderson 2003) using two previous fossil calibrations described in cone snails (Duda Jr. et al. 2001). We excluded *Californiconus californicus* from the CAFE analysis due to optimization failures.

### Diet and conotoxin gene superfamily size evolution

To examine the role of diet specificity and dietary breadth on conotoxin gene superfamily size evolution and total conotoxin diversity, we retrieved prey information from the literature (Kohn 1959a; b, 1966, 1968, 1978, 1981, 2001, 2003, 2015; Marsh 1971; Kohn & Nybakken 1975; Taylor 1978, 1984, 1986; Taylor & Reid 1984; Nybakken & Perron 1988; Kohn & Almasi 1993; Reichelt & Kohn 1995; Kohn et al. 2005; Nybakken 2009; Chang et al. 2015). For diet specificity, we classified prey items into one of 27 different prey categories (Table S9). For dietary breadth, we retrieved estimates of the Shannon’s diversity index (H’) or calculated it if there were at least 5 prey items classified to genus with a unique species identifier. When multiple H’ values were obtained for a species, we averaged them because species will consume different sets of prey taxa depending on geography. Raw data are available in Table S9. To examine the impact of prey group and dietary breadth on changes in gene superfamily size, we used D-PGLS (distance-based phylogenetic generalized least squares), a phylogenetic regression method capable of assessing patterns in high-dimensional datasets (Adams 2014). To reduce redundancy among prey group variables, we removed variables that were 80% correlated with each other using the redun function in the R package Hmisc. We used the total number of exons containing the signal region as our estimate of gene superfamily size. To convert gene superfamily size counts into continuous variables, we transformed the data into chi-squared distances between species in ‘conotoxin gene superfamily space’ using the deostand function in the R package vegan (Oksanen et al. 2016). To examine the impact of diet specificity and dietary breadth on total conotoxin diversity, we used a PGLS analysis implemented in the caper package within R (Orme 2013). We ln-transformed total conotoxin diversity for the PGLS analysis We performed all analyses with the full dataset and a subset of the data that only included gene superfamilies with > 80% capture success. We did not perform any analyses with *C. californicus* because it is regarded as an outlier species amongst the cone snails due to its atypical diet and its deep relationship with the rest of Conidae (Kohn 1966; Puillandre et al. 2014).

## Acknowledgements

We thank DST Hariyanto, MBAP Putra, MKAA Putra, and the staff at the Indonesian Biodiversity Research Center in Denpasar, Bali for assistance in the field in Indonesia; F Criscione, F Köhler, A Moussalli, A Hogget, and L Vail for assistance in the field at the Lizard Island Research Station in Australia; M Reed, A Hallan, and J Waterhouse for access to specimens at the Australian Museum in Sydney, Australia; J Finn, M Mackenzie, and M Winterhoff for access to specimens at the Museum Victoria in Melbourne, Australia; K Bi, L Smith, and A Moussalli for advice on bait design; the Evolutionary Genetics Lab at UC Berkeley for laboratory support; MCW Lim and J Chang for thoughtful advice and discussions; N Puillandre, S Prost, and S Robinson for insightful comments on earlier versions of this manuscript. This work used the Extreme Science and Engineering Discovery Environment (XSEDE), which is supported by National Science Foundation grant number ACI-1053575. This work was supported by a Grants-in-Aid of research from Sigma Xi, a Grants-in-Aid of Research from the Society for Integrative and Comparative Biology, a research grant from the Society of Systematic Biologists, a Student Research Award from the American Society of Naturalists, a National Science Foundation Graduate Research Opportunities Worldwide to Australia, the Lerner Gray Fund for Marine Research from the American Museum of Natural History, research grants from the Department of Ecology and Evolutionary Biology at UCLA, a small award from the B Shaffer Lab, a National Science Foundation Graduate Research Fellowship, an Edwin W. Pauley fellowship, and a Chateaubriand fellowship awarded to MAP. This work used the Vincent J. Coates Genomics Sequencing Laboratory at UC Berkeley, supported by NIH S10 Instrumentation Grants S10RR029668 and S10RR027303. We thank the Indonesian Ministry of State for Research and Technology (RISTEK, permit number 277/SIP/FRP/SM/VIII/ 2013) for providing permission to conduct fieldwork in Bali. The C. californicus specimen was collected under a California Department of Fish and Wildlife collecting permit granted to WF Gilly (SC-6426).

---

**Figure S1. Conotoxin diversity per gene family per region in a phylogenetic context.**

**Figure S2. Scatterplots of all gene superfamilies showing the relationship between exon divergence and noncoding divergence.** Divergence was estimated by calculating uncorrected pairwise distances.

**Figure S3. CAFE net gene gains and losses plotted across a time-calibrated phylogeny of cone snails.** Values on branches represent the number of conotoxin genes gained or loss along that branch. Total conotoxin diversity listed next to species names.

